# Urinary Glycosaminoglycans are Associated with Recurrent UTI and Urobiome Ecology in Postmenopausal Women

**DOI:** 10.1101/2023.01.11.523678

**Authors:** Michael L. Neugent, Neha V. Hulyalkar, Ashwani Kumar, Chao Xing, Philippe E. Zimmern, Vladimir Shulaev, Nicole J. De Nisco

## Abstract

Glycosaminoglycans (GAGs) are linear, negatively charged polysaccharides composed of repeating disaccharide units of uronic acid and amino sugars. The luminal surface of the bladder epithelium is coated with a GAG layer. These urothelial GAGs are thought to provide a protective barrier and serve as a potential interaction site with the urinary microbiome (urobiome). Previous studies have profiled urinary GAG composition in mixed cohorts, but the urinary GAG composition in postmenopausal women remains undefined. To investigate the relationship between GAGs and recurrent UTI (rUTI), we profiled urinary GAGs in a controlled cohort of postmenopausal women. We found that chondroitin sulfate (CS) is the major urinary GAG in postmenopausal women and that urinary CS was elevated in women with active rUTI. We also associated urinary GAGs with urobiome composition and identified bacterial species that significantly associated with urinary GAG concentration. *Corynebacterium amycolatum, Porphyromonas somerae*, and *Staphylococcus pasteuri* were positively associated with heparin sulfate or hyaluronic acid and bacterial species associated with vaginal dysbiosis were negatively correlated to urinary CS. Altogether, this work defines changes in urinary GAG composition associated with rUTI and identifies new associations between urinary GAGs and the urobiome that may play a role in rUTI pathobiology.

## Introduction

Recurrent urinary tract infection (rUTI) is a debilitating disease defined as ≥2 symptomatic urinary tract infections (UTIs) in 6 months or ≥3 UTIs in 12 months.^1-3^ During UTI, uropathogens, such as uropathogenic *Escherichia coli* (UPEC) or *Klebsiella pneumoniae*, ascend the urethra and invade the bladder epithelium (urothelium).^4, 5^ One of the first defenses against infection is a luminal layer of glycosaminoglycan (GAG) polysaccharides coating the bladder epithelium.^6, 7^ These luminal polysaccharides are predominantly composed of the GAGs chondroitin sulfate (CS), heparin sulfate (HS), and hyaluronic acid (HA).^8^ Structurally, GAGs are comprised of sequentially bound disaccharide units of glucuronic acid (or iduronic acid) and amino sugars, such as N-acetylglucosamine (GlcNAc) or N-acetylgalactosamine (GalNAc).^9^ CS and HS are further characterized by sulfated residues (eg., C2, C4, C6 and/or on the non-acetylated nitrogen).^10^ The urothelial GAG layer has been hypothesized to be an important interface between the human host and the resident urinary microbiome (urobiome) or invading uropathogens.^11-13^ For example, Ruggieri et al. reported that acid removal of the GAG layer significantly increased the adherence of *UPEC, Klebsiella ozonae, Proteus mirabilis, Pseudomonas aeruginosa* and *Enterococcus faecalis* in a rabbit model of UTI.^14^

Interestingly, while UPEC is unable to degrade GAGs, the uropathogenic bacterium *Proteus mirabilis* degrades CS, suggesting that some uropathogens may utilize the urothelial GAG layer during urinary tract colonization and invasion.^15^ Additionally, bacterial genera often observed in the urinary microbiome (urobiome), like *Bacteroides* and *Streptococcus* are known to harbor the ability to metabolize GAGs like heparin and HA into smaller disaccharides.^16-20^ Although species of the most predominant urobiome genus, *Lactobacillus*, cannot degrade CS, HS, or HA, *Lactobacillus crispatus, Lactobacillus salivaris*, and *Lactobacillus reuteri* have been shown to adhere to HeLa cells in a GAG-dependent manner.^15, 18, 21^ These observations suggest that the urothelial GAG layer may serve as a scaffolding site for urobiome species.

Aging and menopause, which are also associated with increased UTI susceptibility, may alter the composition of the urothelial GAG layer. Anand et al. reported that estrogen may play a role in modulating GAG layer thickness and the expression of GAG biosynthetic enzymes.^22^ Imamov et al. reported in a female interstitial cystitis mouse model that the quantity of urinary GAGs was higher in estrogen receptor-β knock-out (Erβ^*-/-*^) versus wild-type mice, suggestive of bladder atrophy and urothelium shedding^23^. Deus et al. observed that hypoestrogenism resulted in a lower sulfated GAG content in rat bladders.^24^ Furthermore, GAGs like HA have been shown to be as efficacious as vaginal estrogen for the treatment of symptoms of vaginal atrophy and dyspareunia associated with the decline of systemic estrogen levels in postmenopausal women.^25, 26^

Recently, Bratulic et al. reported reference intervals for the free GAG concentration and disaccharide composition in the urine and plasma samples of 308 healthy adults using a high throughput ultra-high-performance liquid chromatography mass spectrometry (UHPLC-MS/MS) method. In this study, the authors observed higher urinary CS and HS disaccharide concentrations in males than in females.^27^ This work also investigated differences in urinary GAG composition between binned age groups of mixed biological sex and did not find a significant association between serum and urinary GAG concentration and age. Conversely, Larking et al. reported that mean urine creatine-normalized total GAG concentration was bimodal in women and was elevated in postmenopausal compared to premenopausal women.^28^ However, the composition of urinary GAGs in postmenopausal women in the context of rUTI and the organization of the urobiome is poorly understood.

Because rUTI disproportionally affects postmenopausal women, we sought to quantitatively measure urinary GAG composition in a controlled, cross-sectional cohort of postmenopausal women designed to capture the cyclic nature of rUTI using a targeted UHPLC MS/MS-based disaccharide analysis.^8, 29^ We observed that CS was the most abundant urinary GAG in postmenopausal women and that the creatinine-normalized urinary CS concentration was higher in women who were experiencing an active rUTI compared to those not experiencing UTI. We further found evidence that a history of rUTI and estrogen hormone therapy use are associated with altered sulfonation patterns in urinary GAGs. Using whole genome metagenomic sequencing on the same cohort of women, we recently described changes in both the taxonomic composition and functional potential of the urobiome associated with rUTI.^29^ We used this previously described metagenomic dataset to identify urobiome bacterial species associated with urinary GAG concentration. We found that the abundance of taxa associated with urogenital dysbiosis, such as *Atopobium vaginae* and *Peptoniphilus*, negatively correlated with urinary CS concentration.

Conversely, *Corynebacterium amycolatum, Porphyromonas somerae*, and *Staphylococcus pasteuri* were positively associated with HS or HA. Together, our results suggest potential associations between urinary GAG composition and rUTI, estrogen hormone therapy, and urobiome composition.

## Results

### Human cohort design and clinical characteristics

To model the pathobiology of rUTI, we previously curated a controlled cross-sectional cohort of postmenopausal women.^29^ Briefly, postmenopausal women (aged 51-88) were recruited into one of three cohort groups depending on history of rUTI and current UTI status. Group 1 (No UTI History, *n*=25) served as a healthy comparator and consisted of postmenopausal women with no lifetime history of symptomatic UTI. Groups 2 (rUTI History, UTI(-), *n*=25) and 3 (rUTI History, UTI(+), *n*=25) both consisted of postmenopausal women who had a recent history of rUTI but differed in their current status of UTI – Group 2 did not have a symptomatic UTI at the time of sample of collection while Group 3 had a culture-proven symptomatic UTI. To control for common comorbidities that may affect urinary GAG composition or urobiome structure, all postmenopausal women recruited into the cohort passed a strict set of exclusion criteria.^29^

### Targeted liquid chromatography-mass spectrometry assay for urinary GAGs

To quantitatively measure the urinary GAG disaccharide composition of the assembled postmenopausal cohort, we used a modification of previously published targeted liquid chromatography mass spectrometry (LC-MS/MS) methods for the detection of Chondroitin sulfate (CS), Heparin Sulfate (HS), and Hyaluronic Acid (HA) in human urine^8^. All samples were spiked with 100ng of a non-naturally occurring heparin disaccharide. Derivatized samples were analyzed via targeted LC-MS/MS on a Waters Xevo TQS and ACQUITY UPLC using multiple reaction monitoring (MRM) for parent>daughter transitions representing the 17 disaccharides of interest from CS, HS, and HA (Figure 1 A & B, Table 1, Supplemental Figure 1). We obtained robust and reproducible assay performance over a linear analytical range from 10ng/mL to 1000ng/mL for all analytes of interest (Supplemental Figure 1).

**Figure 1.**
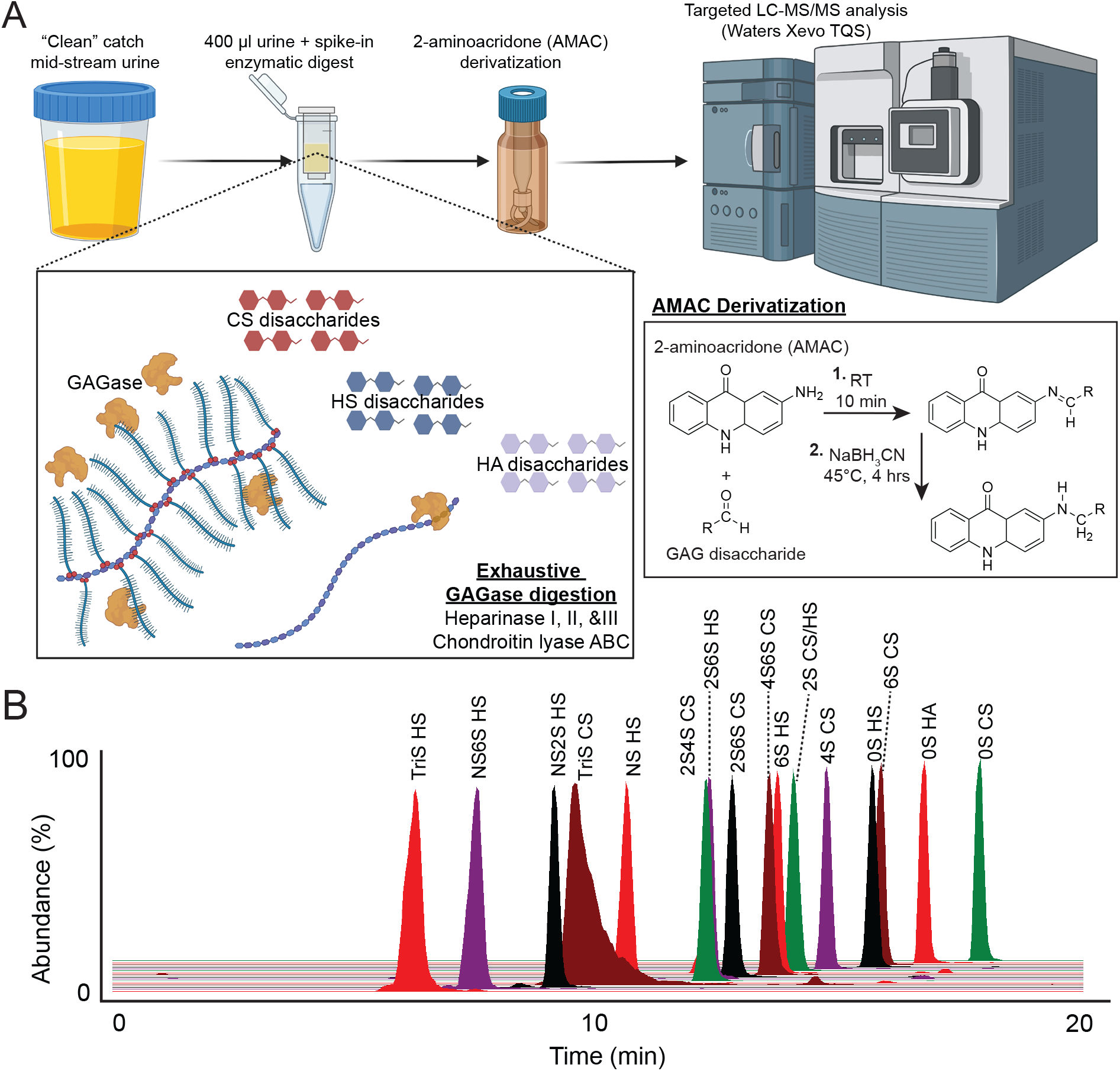
Targeted LC-MS/MS assay for urinary GAGs CS, HS, and HA. (A) Schematic diagram of LC-MS/MS assay workflow from sample preparation, enzymatic digestion, derivatization, and analysis by LC-tandem quadrupole mass spectrometer. (B) Chromatogram of eluting GAG disaccharide analytical MRM transitions.

### All three major GAGs are detectable in postmenopausal urine, but CS predominates

We observed all 17 expected disaccharides for CS, HS, and HA.^8, 30^ The most abundant disaccharides were the mono-sulfate and non-sulfate CS disaccharides 4S-CS (2460.5 ng/mL, CI95%=2262-3193 ng/mL), 6S-CS (1524 ng/mL CI95%=1192-1945 ng/mL), and 0S-CS (1472 ng/mL CI_95%_=1219-1827 ng/mL) (Figure 1 2A & B). We also observed abundant levels of the non-sulfate HS disaccharide, 0S-HS (505.7 ng/mL CI_95%_=374.1-648.4 ng/mL), the HA disaccharide, 0S-HA (592.2 ng/mL CI_95%_=428.2-716.7 ng/mL), and the di-sulfate 4S6S disaccharide of CS (236.6 ng/ml CI_95%_=188.1-283.7 ng/mL) (Figure 2A & B, Supplemental Table 1 & 2).

**Figure 2.**
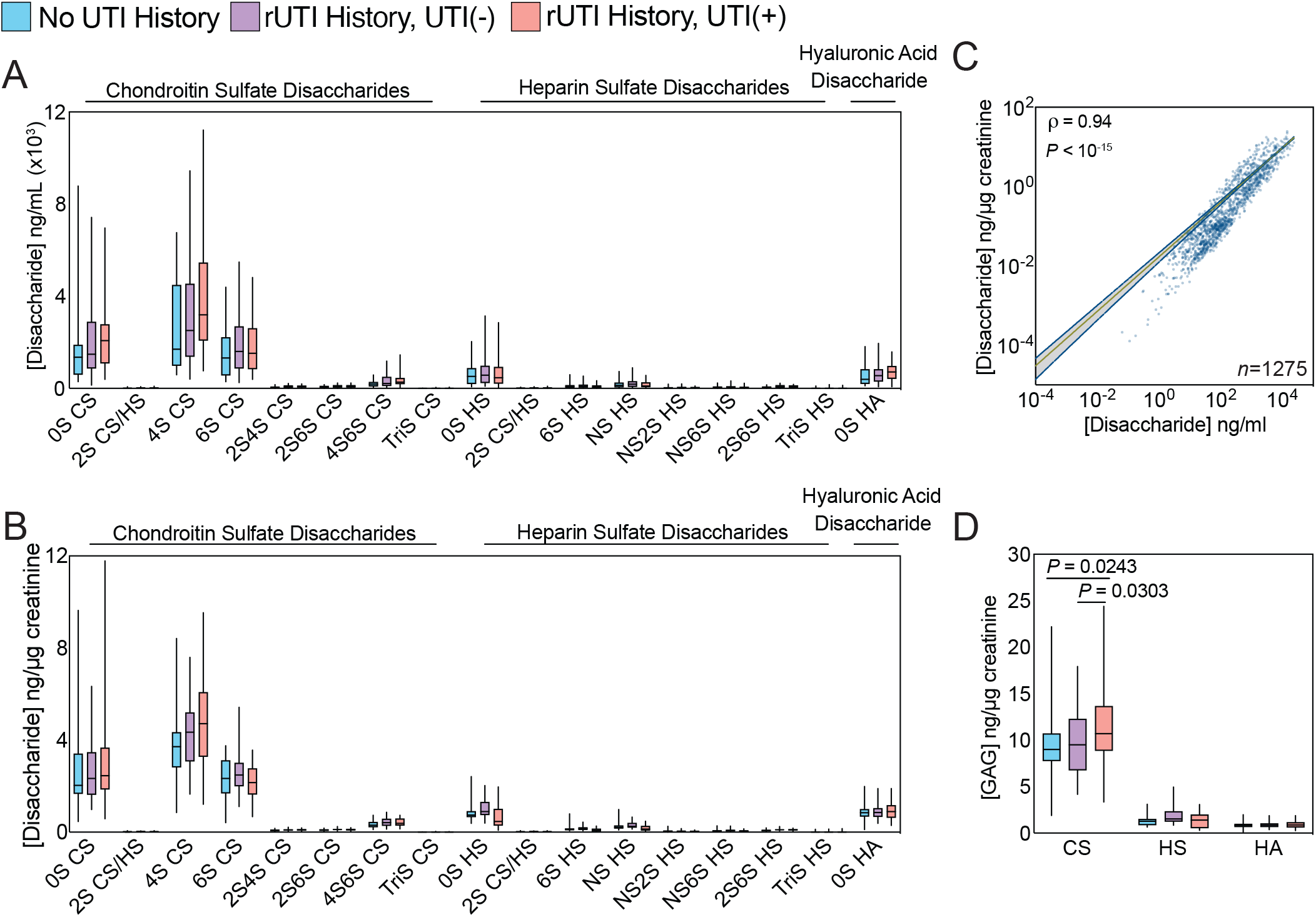
Urinary GAG profile in postmenopausal women with and different UTI histories. (A) Urinary GAG disaccharide concentration in ng/mL. (B) Urinary GAG disaccharide concentration normalized to urinary creatinine. (C) Correlation of normalized and unnormalized urinary GAG disaccharide concentrations. P-value generated by t-ratio calculation. (D) Comparison of summed GAG disaccharide distributions across cohort groups (n=25 in each group, n=75 total). P-value generated by 2-way ANOVA with Fischer LSD multiple comparison post-hoc. All boxes are drawn to represent the interquartile range. Median is denoted by solid horizontal line. Bars are drawn from minimum to maximum of the data range.

To account for hydration status (i.e. urine concentration), urinary metabolites are often normalized to urinary creatinine.^31^ However, it has been suggested that urinary creatinine excretion rate may not be constant between and within individuals.^32^ To assess the agreement between raw urinary disaccharide concentration and creatinine-normalized disaccharide concentration, we performed linear regression and correlation analysis between the raw and creatinine-normalized datasets. We observed a strong linear correlation (Spearman ρ=0.94, *P*<10^−15^) when comparing raw disaccharide concentrations with creatinine-normalized abundance (Figure 2 C). We chose to use creatinine-normalized data for downstream analyses due to convention on how urinary metabolites are often reported and the strong linear association we observe between raw and creatinine-normalized concentrations (Supplemental Table 2).

Consistent with previous observations, our analysis found that the most abundant GAG detected in postmenopausal urine was CS (Figure 2D).^8^ Interestingly, we observed significantly higher levels of urinary CS in urines from patients with active rUTI (Figure 2D). This observation was most significant when comparing creatinine-normalized urinary CS concentration between the No UTI History and rUTI History, UTI(+) groups (*P*=0.0243) and may be indicative of tissue damage and urothelial barrier disruption occurring during UTI.^11, 33^ No difference was found in total urinary HS or HA concentration between cohort groups.

### rUTI history is associated with altered urinary GAG sulfonation pattern

Given that we observed a significant positive association between urinary CS concentration and active rUTI, we hypothesized that the sulfonation pattern of urinary GAGs may differ between women with different rUTI histories. To test this hypothesis, we performed a pairwise metabolite enrichment analysis between the groups in our cohort using the non-parametric Wilcoxon rank sum test. When comparing the No UTI History group with the rUTI History, UTI(-) group, we detected the enrichment of disaccharides 2S6S-HS (*P*=0.0037) and 4S6S-CS (*P*=0.0353) in the rUTI History, UTI(-) group (Figure 3 A-C). Similarly we observed enrichment of the 2S6S-HS disaccharide in the rUTI History, UTI(+) group compared to the No UTI history group (*P*=0.0080) (Figure 3 D, E). We also observed an elevation of the 2S4S-CS disaccharide in the rUTI History, UTI(+) group urine (*P*=0.0472) and an elevation of the mono-sulfonated 6S-HS disaccharide in No UTI History group urine (*P*=0.0450) (Figure 3 F, G). These results suggest that rUTI history may be associated with remodeling of urinary GAG sulfonation patterns of HS and CS. Our analysis further detected significant enrichment of HS disaccharides in the rUTI History, UTI(-) group compared to the rUTI History, UTI(+) group (Supp Figure 2 A). These differential features included the enrichment of mono-sulfate 6S- and NS-HS disaccharides (*P*=0.0016, *P*=0.0034), the di-sulfate NS6S- and NS2S-HS disaccharides (*P*=0.0102, *P*=0.0353), and the non-sulfate 0S-HS disaccharide (*P*=0.0161) in the rUTI History, UTI(-) group (Supp Figure B-F). Taken together, these observations provide rationale to hypothesize that rUTI history changes the composition of urinary GAG sulfonation patterns in postmenopausal women.

**Figure 3.**
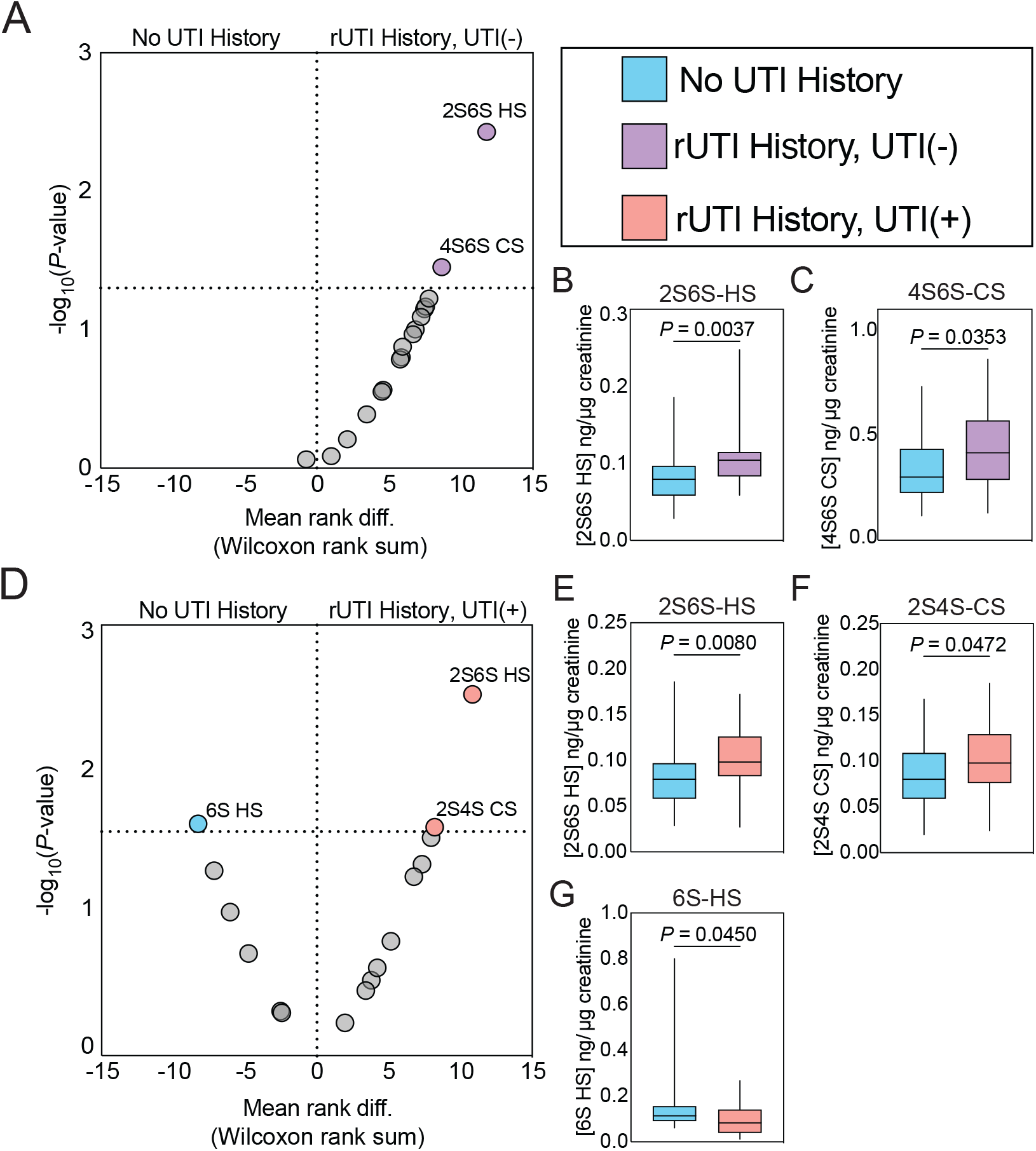
rUTI history is associated with altered urinary GAG disaccharide composition. (A) Volcano plot depicting differential enrichment screening of urinary GAG disaccharides between the No UTI History (n=25) and rUTI History, UTI(-) groups (n=25). (B, C) Box plots depicting significant hits from differential enrichment screen. (D) Volcano plot depicting differential enrichment screening of urinary GAG disaccharides between the No UTI History (n=25) and rUTI History, UTI(+) groups (n=25). (E, F, G) Box plots depicting significant hits from differential enrichment screen. All P-values generated by the non-parametric Wilcoxon rank-sum. All boxes are drawn to represent the interquartile range. Median is denoted by solid horizontal line. Bars are drawn from minimum to maximum of the data range.

### Estrogen hormone therapy is associated with altered urinary CS sulfonation

Many postmenopausal women use estrogen hormone therapy (EHT) to alleviate symptoms of menopausal syndrome.^34^ We previously showed with this cohort that EHT strongly shapes the taxonomic profile of the postmenopausal urobiome, enriching for protective species, such as *Lactobacillus crispatus*.^29^ Interestingly, previous reports also suggest that EHT thickens the luminal GAG layer of the urothelium and regulates the GAG sulfotransferase enzyme HS6ST1.^22, 35^ Given these observations, we hypothesized that EHT may augment urinary GAGs derived from the urothelium. To test this hypothesis, we compared total urinary CS, HS, and HA abundance among the No UTI History and rUTI History, UTI(-) groups now dichotomized by EHT use. We observed no difference in total urinary CS, HS, or HA abundance (Figure 4A). We further performed a metabolite enrichment analysis of urinary GAG disaccharides among the EHT(+) and EHT(-) groups using the non-parametric Wilcoxon rank sum test (Figure 4B). This analysis identified two urinary CS disaccharides, 4S6S-CS and 2S-CS/HS, as significantly enriched in the EHT(+) patients (*P*=0.0174 and *P*=0.0346, respectively) (Figure 4 C & D). Given the inherent difficulty to distinguish the 2S-CS and 2S-HS disaccharides owing to their identical MRM transition and similar chromatographic retention, we grouped the feature as an aggregate of the two possible disaccharides. While we did not find evidence to support our hypothesis that EHT augments the total amount of urinary GAGs, these data do suggest that EHT may alter the sulfonation patterns of urinary GAGs.

**Figure 4.**
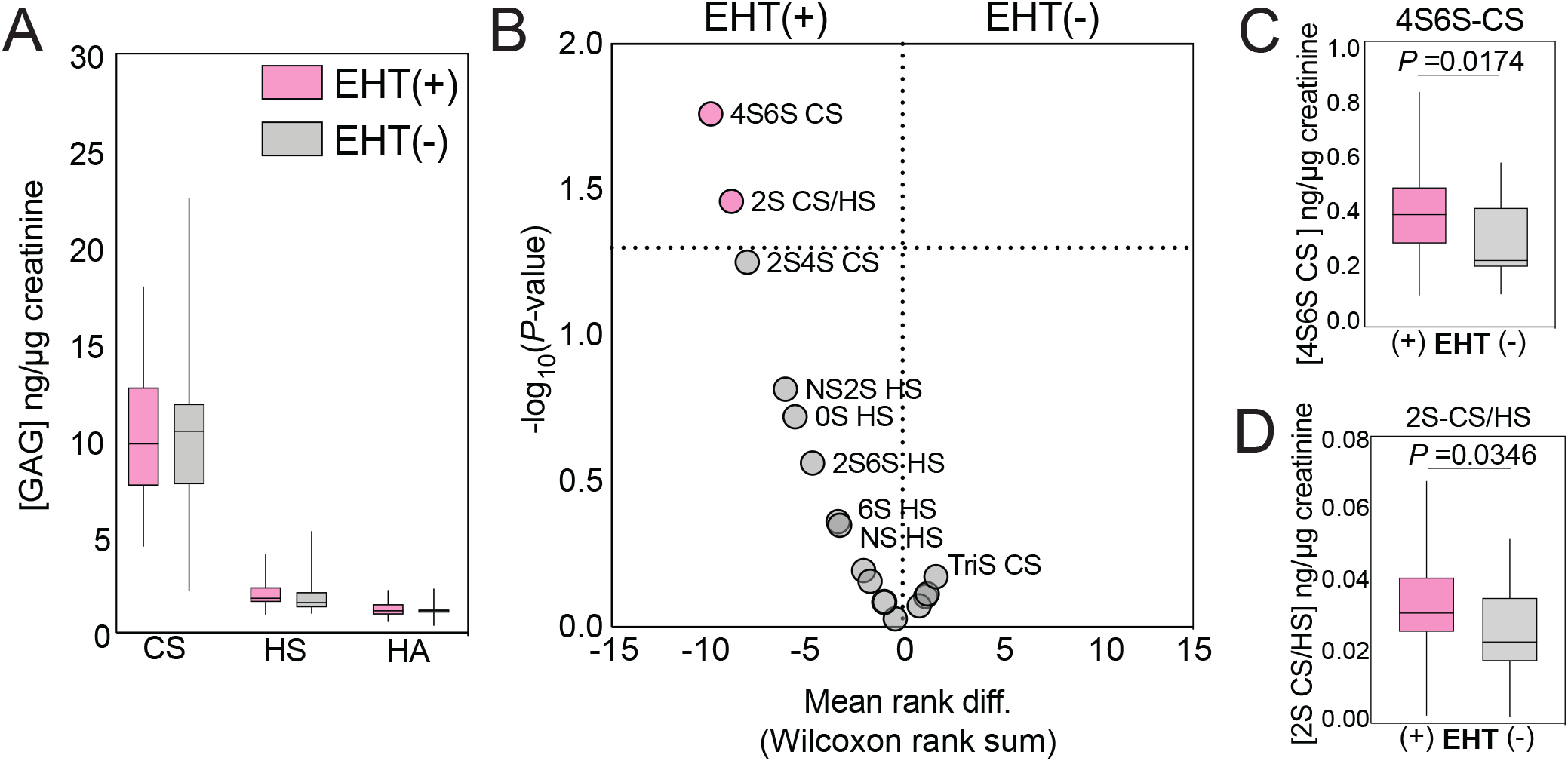
Estrogen hormone therapy is associated with altered urinary CS disaccharide composition. (A) Comparison of summed GAG disaccharide distributions between EHT(+) (n=29) and EHT(-) (n=21) women on the pooled No UTI History and rUTI History, UTI(-). (B) Volcano plot depicting differential enrichment screening of urinary GAG disaccharides between the between EHT(+) (n=29) and EHT(-) (n=21) women. (C, D). Box plots depicting significant hits from differential enrichment screen. All P-values generated by the non-parametric Wilcoxon rank-sum. All boxes are drawn to represent the interquartile range. Median is denoted by solid horizontal line. Bars are drawn from minimum to maximum of the data range.

### Urobiome bacterial species associated with urinary GAGs

The urinary tract hosts a unique microbial ecosystem, members of which belong to the urobiome.^4, 36-43^ A decade of work has studied and defined the urobiome in multiple diseases, including overactive bladder, bladder cancer, urinary incontinence, and rUTI.^4, 29, 44-46^ However, host factors contributing to the urobiome ecosystem are poorly understood. We previously performed whole genome metagenomic sequencing of the urobiome in the cohort studied here and reported differential taxonomic and functional characteristics associated with rUTI in postmenopausal women.^29^ To assess urobiome bacterial ecology associated with urinary GAG composition, we performed a correlative screen of bacterial species with urinary GAG concentration in matched samples. Using the non-parametric Spearman correlation (ρ), we identified five associations between urinary GAGs and urobiome bacterial species that passed both a threshold of statistical significance (*P*<0.05) and correlation strength (|ρ| ≥ 0.3) (Supplemental Table 3). Three positive correlations were identified between HS and *Corynebacterium amycolatum* (ρ= 0.345, *P*=0.0024), HA and *Porphyromonas somerae* (ρ= 0.319, *P*=0.0053), and HS and *Staphylococcus pasteuri* (ρ= 0.309, *P*=0.0071) (Figure 5A). We further observed two significant negative correlations between CS and *Atopobium vaginae* (ρ= -0.363, *P*=0.0014) and CS and *Peptoniphilus rhinitidis* (ρ= -0.316, *P*=0.0057). To assess these associations further, we dichotomized the cohort population by presence or absence of these identified bacterial species and compared GAG abundance between (+) and (-) groups. We observed significantly higher urinary HS in patients with *C. amycolatum* or *S. pasteuri* (*P*=0.0016, *P*=0.0045) as well as significantly higher urinary HA in patients with *P. somerae* (*P*=0.003) (Figure 5 B, C, D). Significantly lower urinary CS concentrations were observed in patients with *A. vaginae, P. rhinitidis* detected in their urobiome (*P*=0.0012, *P*=0.0068). While not meeting the strict correlation cutoff of |ρ| > 0.3 of our initial correlative screen (ρ= -0.282, *P*=0.0142), we did observe significantly lower urinary CS concentrations in patients with *Ruminococcus obeum* (renamed *Blautia obeum*) present in their urobiome (*P*=0.0154) (Figure 5 G). Taken together, these data suggest that urinary GAGs associate with the urobiome and thus, may play a role in shaping the bacterial ecology of this environment.

**Figure 5.**
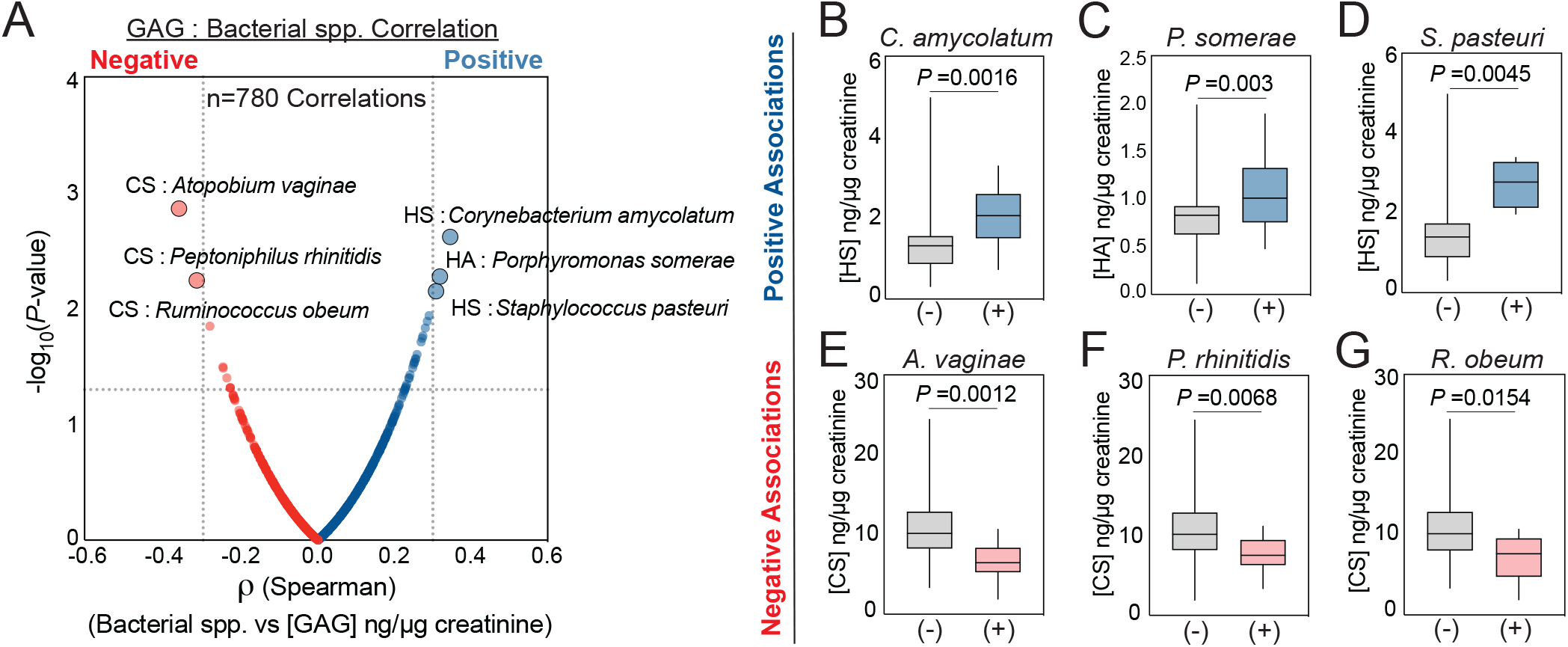
Urobiome bacterial species are associated with urinary GAG composition. (A) Volcano plot depicting correlation analysis of summed urinary GAGs with urobiome bacterial species detected by whole genome metagenomic sequencing (n=260 species). For the Spearman correlation, thresholds of ρ ≥ 0.3 and *P* < 0.05 were chosen to assign significance. P-value generated by t-ratio calculation. Positive GAG-Bacteria correlations are colored blue, negative are colored red. (B, C, D) Box plots depicting significant positive hits from correlation analysis screen. Patients are dichotomized by presence(+)/absence(-) of identified species as follows: *C. amycolatum* n(-)=60, n(+)=15; *P. somerae* n(-)=47, n(+)=28; *S. pasteuri* n(-)=71, n(+)=4. (E, F, G) Box plots depicting significant negative hits from correlation analysis screen. Patients have been dichotomized by presence(+)/absence(-) of identified species as follows: *A. vaginae* n(-)=66, n(+)=9; *P. rhinitidis* n(-)=63, n(+)=12; *R. obeum* n(-)=67, n(+)=8. P-values depicted in boxplots were generated by the non-parametric Wilcoxon rank-sum. All boxes are drawn to represent the interquartile range. Median is denoted by solid horizontal line. Bars are drawn from minimum to maximum of the data range.

## Discussion

rUTI is a public health problem disproportionately affecting postmenopausal women who currently are underrepresented in biomedical science.^4, 29^ In this population, rUTI can persist for years leading to a dramatic reduction in quality of life, massive financial burden, and life-threatening infections if treatment is unsuccessful.^1^ Yet, little is known about the pathobiology of rUTI particularly in postmenopausal women. Understanding the urogenital environment is critical to further our understanding of rUTI susceptibility and to develop new therapies. An important defense mechanism and site of host-pathogen or host-microbiome interaction in the urinary tract is the luminal urothelial GAG layer.^6, 7, 35^ It is believed that the female urobiome, which has now been identified as a key component of the urogenital environment, interacts with the urothelial GAG layer.^36-43^ However, little is currently known about how the urobiome as a whole and how individual urobiome species associate with the urothelial GAG layer or urinary GAG composition. Here, we used targeted mass spectrometry-based metabolite profiling to quantitatively measure urinary GAGs in a cohort of postmenopausal women designed to study rUTI. Our observations that CS is the major urinary GAG in this cohort is consistent with previous findings and consistent with the composition of the luminal GAG layer of the urothelium.^8^ We further observe that rUTI alters not only the total amount of urinary GAGs but also the sulfonation patterns of urinary GAGs. These observations may suggest that active rUTI may disrupt the urothelial GAG layer integrity and sulfonation. These data provide rationale to mechanistically study the urothelial GAG layer in rUTI pathobiology. Future work will focus on assessing the transitional and biological relevance of elevated urinary CS during active rUTI as well as the longitudinal stability of these observations. Interestingly, while we observed an association between urinary CS sulfonation and EHT use, we did not observe differences in total GAG abundance between women using and not using EHT. It should be noted that a limitation of this study is that we assayed free urinary GAGs which may not be fully reflective of urothelial GAG layer thickness, integrity, or composition. Tissue samples would likely be needed to robustly determine if EHT is associated with specific augmentations of the thickness or composition of the urothelial GAG layer. We did, however, observe that EHT use was associated with altered urinary GAG sulfonated disaccharide patterns. We interpret these findings as supportive of previous work described by Anand et al. that showed differential expression of GAG sulfotransferase and biosynthesis enzymes between ovariectomized and Sham mice.^22^

A body of evidence suggests that both EHT and intravesical administration of GAGs have the potential to reduce the incidence of UTI and rUTI in human subjects.^47-50^ Ablove et al. reported that intravesical administration of HA and heparin prevented recurrent bacterial cystitis and refractory rUTI respectively.^51, 52^ Additionally, HA-CS intravesical instillations were found to significantly reduce the incidence of rUTI in women with a history of rUTI.^53^ These findings are supported by studies in rat models of cystitis that demonstrate a decrease in bladder *E. coli* burden after HA instillation. Moreover, this work demonstrated that instilled HA was able to coat the urothelium and integrate into the GAG layer, suggesting that therapeutically delivered GAGs may increase barrier function and protect against invasion of *E. coli* into the urothelium.^54^ Taken together, these data beg the question of whether EHT and/or intravesically delivered GAGs can be used to therapeutically augment the urothelial GAG layer to treat urological diseases in postmenopausal women.

The urobiome is another potential target for new rUTI therapeutics.^4^ This understudied microbial environment may harbor intricate interactions between specific host factors and members of the urobiome. However, our knowledge about the interaction of the urobiome with the urothelium and urinary GAGs is limited. Association of *Lactobacillus* species *Lactobacillus crispatus, Lactobacillus salivarius, Lactobacillus reuteri* with HeLa cells has been reported to be GAG-dependent.^18, 21^ Interestingly, depletion of the GAG layer in the bladder of a rabbit UTI model led to an increased attachment of UPEC and *Klebsiella pneumoniae*, suggesting that the GAG layer may impede uropathogen adherence to the urothelium.^13, 55, 56^ Indeed, it is known that some uropathogens, such as *Streptococcus agalactiae* and *Proteus mirabilis*, can degrade CS.^15^ We find it interesting to note that all the negative taxa-GAG associations were observed with CS, the most abundant urinary GAG.^8^ Also of note, the presence of *A. vaginae* and species of *Peptoniphilus*, which were negatively associated with urinary CS, are known to be a signature of dysbiosis in the vagina.^29, 57-62^ Future work will need to investigate mechanistic links between negative associations we observed between *A. vaginae, Peptoniphilus*, and urinary CS in order to determine if these taxa directly impact or interact with GAGs or are instead part of larger signature of dysbiosis and dysfunction in the urinary tract.

## Conclusion

Urinary GAGs are associated with rUTI disease state and urobiome ecology. We hypothesize that urogenital dysbiosis is also associated with changes in the urothelial GAG layer. This hypothesis is evidenced by our observations of negative correlations between urinary CS and taxa known to be signatures of dysbiosis in the vagina. The work presented here provides strong rationale to mechanistically study the impact of the urothelial GAG layer on urobiome composition and structure.

## Acknowledgements

The authors would like to thank and acknowledge the patients who participated in this study. This work was supported by grants from the Welch Foundation (AT-2030020200401) and the National Institutes of Health (1R01DK131267-01) to N.J.D and (1F32DK128975-01A1) to M.L.N. This work was also supported by the Felicia and John Cain Distinguished Chair in Women’s Health to P.E.Z.

## Author Contributions

Conceptualization, M.L.N., N.H., V.S., N.J.D; data curation, M.L.N., N.H., A.K., C.X., P.E.Z., V.S., N.J.D; formal analysis, M.L.N., N.H., V.S., N.J.D.; funding acquisition, M.L.N., P.E.Z., N.J.D.; investigation, M.L.N., N.H.; methodology, M.L.N., N.H., A.K., C.X., P.E.Z., V.S. N.J.D.; project administration, N.J.D.; resources, P.E.Z., V.S., N.J.D.; supervision, C.X., P.E.Z., V.S., N.J.D.; validation, M.L.N., N.H., V.S., N.J.D; visualization, M.L.N., N.J.D.; writing-original draft, M.L.N., N.H., N.J.D.

### Experimental Section

#### Cohort Curation

The human cohort presented in this study is approved under IRBs STU032016-006 (University of Texas Southwestern Medical Center) and 19MR0011 (University of Texas at Dallas) and was recruited as previously reported.^29^ Briefly, participants were recruited with written, informed consent from a single site in the Urology clinic at the University of Texas Southwestern Medical Center between 2018 and 2019. All participants were postmenopausal females who passed a set of exclusion criteria: pre-or perimenopausal, complicated rUTI (i.e. indwelling catheter), pelvic malignancy, history of pelvic radiation within 3 years, pelvic floor procedure within 6 months prior, ongoing chemotherapy, renal insufficiency (creatinine >1.5 mg/dL), >stage 2 prolapse, Diabetes Mellitus (DM) type 1 or 2, neurogenic bladder, upper urinary tract anomaly, and post void residual (PVR) > 100 mL. Patients were also excluded if antibiotic were used within 4 weeks prior unless a positive urine culture test result was observed. All urine samples were collected via “clean-catch” midstream method after participant education about the requirements for collection. Urine samples were stored at 4°C ≤3hrs before sample processing and storage at -80°C.

#### Urobiome whole genome metagenomics

Urinary whole genome metagenomics was previously reported for the cohort presented here.^29^ Briefly, metagenomic DNA was isolated from pelleted urine samples using the Zymo Research DNA/RNA microbiome miniprep kit. DNA quality was assessed by Qubit assay and agarose gel electrophoresis. DNA libraries were constructed using the Nextera DNA Flex kit. Whole genome metagenomic sequencing was performed at the University of Texas at Dallas Genome Center using 2×150 bp paired-end reads on an Illumina NextSeq500 with a target of ≥50 million paired end reads per sample.

#### Bioinformatic analysis

Bioinformatic analysis of metagenomic data was performed and reported previously.^29^ Taxonomic analysis was performed by first trimming the raw FASTQ files of human-mapping reads with KeadData.^63^ MethaPhlAn2 was used to generate taxonomic profiles.^64^ For complete information detailing how these analyses were performed, we refer the reader to the primary report of these data by Neugent et al.^29^

#### Urine desalting and urinary GAG concentration

We used a modified version of previously reported methods to quantify urinary GAGs.^8^ Urine samples (stored at -80C) were thawed at room temperature. 400 µL of each urine sample was loaded onto a 3 kDa molecule weight cutoff centrifugal filter column unit (Amicon UFC500396) and placed at 37°C water bath for 20 minutes (or until the sample turned clear and devoid of any precipitate). Desalting was carried out by spinning the sample-containing filtration column for 1 hour at 14000xg. The column was washed twice with deionized water for 45 minutes at 14000xg each. Additionally, urine samples spiked with 10 ng/mL of CS and HS each were prepared for method validation as positive controls.

#### Exhaustive enzymatic digestion of urinary GAGs

Following the filter column wash, the casing/collection tubes were replaced prior to digestion. 150 µL digestion buffer (50 mM ammonium acetate containing 2 mM calcium chloride; pH 7.0) was added to the filter column. Recombinant heparin lyase I, II, III (pH optima 7.0-7.5) and recombinant chondroitinase ABC (chondroitin lyase) (pH optimum 7.4) were added to each sample at 10 mU each and mixed well. Enzymatic digestion was performed at 37°C water bath for 2 hours. The digestion process was terminated by centrifugation at 14000xg for 1 hour. The filter column was washed twice with 100 µL deionized water, each for 45 minutes at 14000xg. Deionized water sample was used as a digestion control for the method.

#### Disaccharide Derivatization with 2-aminoacridone

Each sample filtrate containing the disaccharide products was spiked with 100 ng/µL of a non-naturally occurring Heparin Sulfate disaccharide (∆UA-2S GlcNOEt-6S), which has a -COCH2CH3 moiety attached to the amine of the glucosamine, as an internal standard (Iduron United Kingdom). This was followed by drying using a vacuum centrifuge. After drying, 10 µL of 0.1M AMAC in DMSO/acetic acid (17/3, V/V) was added to each sample and incubated at RT for 10 minutes. 10 µL of 1M aqueous sodium cyanoborohydride was added and derivatization was performed at 45°C for 4 hours. The samples were then centrifuged at 14000xg for 5 minutes to recover the supernatant and 60 µL of DMSO: acetic acid: deionized water (17:3:20) was added. Samples were stored in a light-resistant amber vial at 4°C until analyzed by LC-MS/MS.

#### Liquid chromatography mass spectrometry

Quantitative measurement of urinary GAG disaccharides was performed via a modification of previously reported methods on a Waters Xevo TQ tandem quadrupole mass spectrometer, in negative ion mode, lined to and ACQUITY UPLC equipped with a CORTECS UPLC T3 2.1×150 mm UPLC column with a 1.6 µm particle size.^8^. 5 µL of derivatized sample was injected to run on a 31-minute liquid chromatography gradient starting at 95% 80 mM ammonium acetate p.H. 5.8 (solvent A) to 100% methanol (solvent B) at a 0.3mL/min flowrate. Column temperature was held at 40°C. Previously reported MRM libraries of GAG disaccharides were validated from pure standards of the 17 disaccharides of interest (Iduron, United Kingdom) and used to monitor analytical transitions (Table 1).^8^ Data analysis was performed in TargetLynx. Briefly, analytical transition peaks for each disaccharide were integrated, normalized to internal spike-in standard and mapped to a standard curve to accurately estimate analyte concentration. Urinary disaccharide concentrations were then normalized to urinary creatinine, measured by colorimetric assay (Sigma) to account for urine concentration and hydration status.

#### Statistical analysis

Statistical analysis was performed using GraphPad Prism 9 and Microsoft Excel. The non-parametric Wilcoxon rank sum was used for pairwise hypothesis testing. The Spearman correlation was performed as a non-parametric method of assessing the association of two continuous variables. An alpha of 0.05 was used to control type I error.

