## Supplemental Material for "Urinary Glycosaminoglycans are Associated with Recurrent UTI and Urobiome Ecology in Postmenopausal Women"

### Supplemental Figure 1

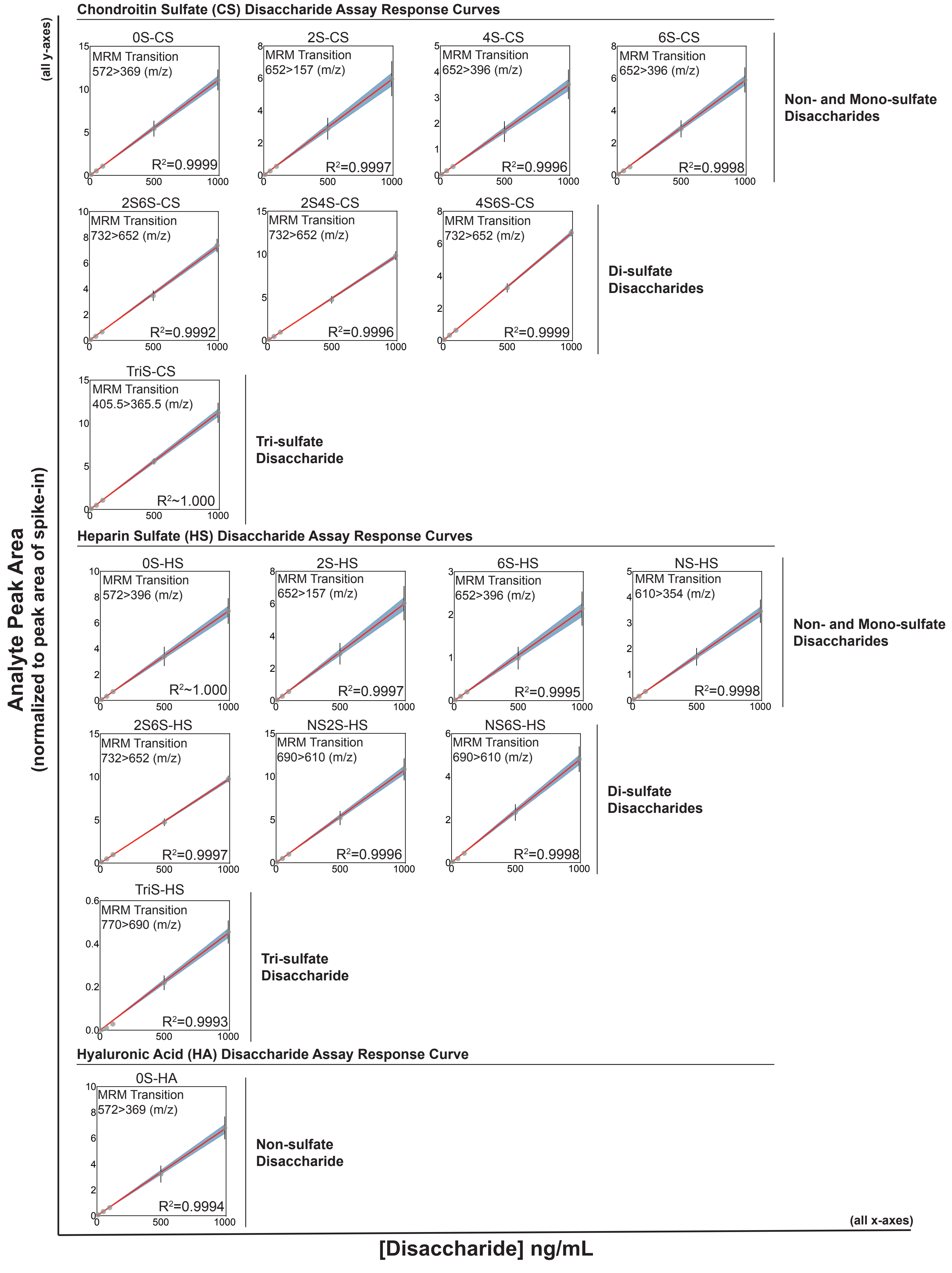

### Supplemental Figure 2

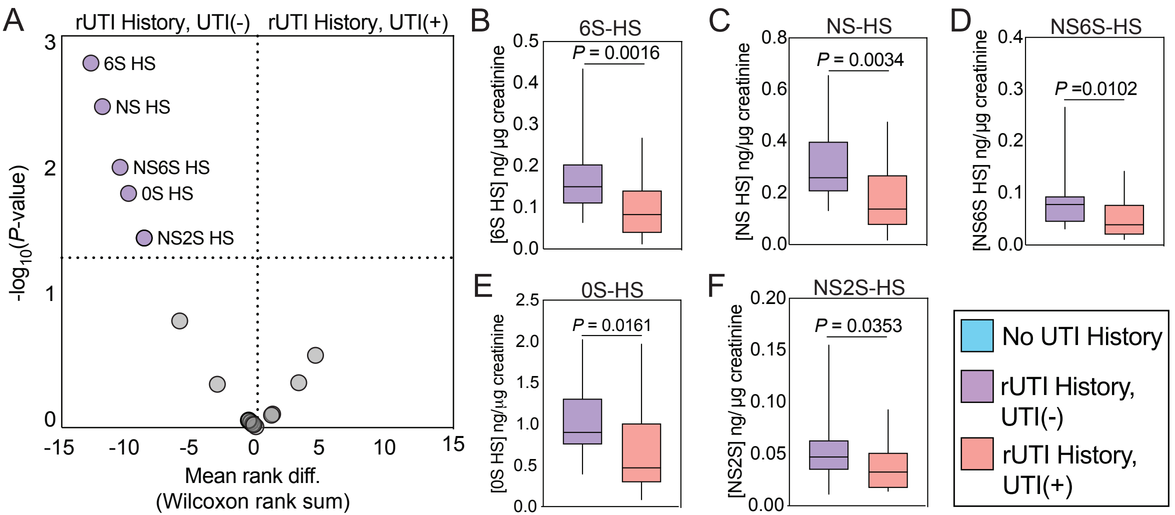

**Supplemental Figure 1. Response curves for all 17 monitored disaccharide MRM analytical transitions.** Scatter plots with linear regression and correlation analysis of all monitored disaccharide MRM analytical transitions for CS (n=8 transitions), HS (n=8 transitions), HA (n=1 transition). All y-axes represent analyte peak area normalized to the peak area of the internal spike-in. All x-axes represent the concentration of disaccharide in ng/mL. $R^2$ generated from the Pearson coefficient. 95% confidence region of the red linear regression line is shaded in blue. Data presented is an aggregate of 3 independent instrument runs performed in different days. Dots represent the mean of replicates. Error bars represent the range (minimum to maximum) of replicate assay response.

**Supplemental Figure 2. Comparison of GAG sulfonation patterns between rUTI History, UTI(-) and rUTI History, UTI(+) groups.** (A) Volcano plot depicting differential enrichment screening of urinary GAG disaccharides between the rUTI History, UTI(-) (n=25) and rUTI History, UTI(+) (n=25) groups. (B-F) Box plots depicting significant hits from differential enrichment screen. All P-values generated by the non-parametric Wilcoxon rank-sum. All boxes are drawn to represent the interquartile range. Median is denoted by solid horizontal line. Bars are drawn from minimum to maximum of the data range.
